# The Establishment of CDK9/ RNA PolII/H3K4me3/DNA Methylation Feedback Promotes HOTAIR Expression by RNA Elongation Enhancement in Cancer

**DOI:** 10.1101/812776

**Authors:** Chi Hin Wong, Chi Han Li, Qifang He, Joanna Hung Man Tong, Ka-Fai To, Yangchao Chen

## Abstract

Long non-coding RNA HOX Transcript Antisense RNA (HOTAIR) is overexpressed in multiple cancers with diverse genetic profiles, which heavily contributed to cancer progression. However, the underlying mechanism leading to HOTAIR deregulation is largely unexplored. Here, we revealed that gene body methylation promoted HOTAIR expression through enhancing the transcription elongation process in cancer. We linked up the aberrant gene body histone and DNA methylation in promoting transcription elongation via phosphorylation of Polymerase II Ser 2 by CDK7-CDK9, and elucidated the mechanism of a positive feedback loop involving CDK7, MLL1 and DNMT3A in promoting gene body methylation and overexpressing HOTAIR. To our knowledge, this is the first time to demonstrate that a positive feedback loop that involved CDK9-mediated phosphorylation of PolII and histone and gene body methylation induced robust transcriptional elongation, which heavily contributed to the upregulation of oncogenic lncRNA in cancer.

## INTRODUCTION

HOX Transcript Antisense RNA (HOTAIR), as one of the well-studied long non-coding RNA (lncRNA), is significantly overexpressed in multiple cancers with diverse genetic profiles, including both solid and hematopoietic cancers (Tang et al., 2018). HOTAIR drives important cancer phenotypes via inducing deregulations on critical genes by PRC2-mediated H3K27 trimethylation, LSD1/CoREST/REST-mediated H3K4 demethylation and interacting with miRNAs as competitive RNA (Li et al., 2017; Liu et al., 2013; Zhang et al., 2015). However, the mechanism in regulating HOTAIR expression in cancer remains unexplored.

Methylation of DNA at cytosine of CpG dinucleotides has been reported to regulate gene expression (Antequera, 2003). A previous report suggested that methylation in downstream CpG island (DS-CGI) of HOTAIR gene facilitated the transcription of HOTAIR in breast cancer (Lu et al., 2012). The DS-CGI, which is located between HOTAIR gene and HOXC12 gene, controls the expression of HOTAIR. When DS-CGI is methylated, transcription of HOXC12 terminated at DS-CGI and HOTAIR can be transcripted normally. However, when DS-CGI is unmethylated, transcription of HOXC12 continues and disrupts HOTAIR transcription. In this study, we attempted to delineate the underlying mechanism leading to the deregulation of HOTAIR.

Here, we found that methylation of DS-CGI may not be involved in regulation of HOTAIR expression. Importantly, we demonstrated a general mechanism that contributed to HOTAIR upregulation in cancers. Methylation of exon CpG island in HOTAIR gene promoted HOTAIR expression by facilitating transcription elongation process. In addition, the methylation of exon CpG island was regulated by CDK9-mediated phosphorylation of RNA PolII and MLL1-mediated H3K4me3.

## RESULTS

### Methylation of downstream CpG island may not be involved in regulating HOTAIR expression

HOTAIR was significantly upregulated in many cancer types and played important roles in cancer progression (Figure S1). However, the mechanism in regulating HOTAIR expression in cancer remains unexplored. Previous report hypothesized that the methylation of CpG island, which is downstream of HOTAIR gene, was required for transcription of full-length HOTAIR transcript (Figure S2A) (Lu et al., 2012). Therefore, we attempted to validate this theory. Firstly, we profiled the HOTAIR expression and the methylation status of DS-CGI in a panel of breast cancer cell lines. We failed to observe the correlation relationship between HOTAIR expression and DS-CGI methylation (Figure S2B, C). Furthermore, both downstream 5.8 kb and upstream 4 kb transcript of the DS-CGI could be detected in all breast cancer cell lines regardless of methylation status (Figure S2C). Regarding to the situation in PDAC, both the 5.8 kb and the 4 kb transcript were not detected in non-tumor HPDE cells with low HOTAIR level (Figure S2D). Chromatin immunoprecipitation (ChIP) analysis also demonstrated that RNA Polymerase II enriched at 4 kb transcript, suggesting active transcription of the 4 kb transcript in SW1990 cells with high HOTAIR level (Figure S2D). These findings contradicted to the previous theory that methylation of DS-CGI hindered the transcription of downstream 4 kb transcript. Therefore, we here demonstrated that methylation of DS-CGI may not be involved in regulating HOTAIR expression.

### DNA methylation in Exon-CpG Island contributed to HOTAIR expression in cancer

To study the association between DNA methylation and HOTAIR expression, we located all CGIs in the HOTAIR gene region. Apart from DS-CGI which was not involved in regulating HOTAIR expression, we observed another two CGIs in HOTAIR gene (Figure 1A). We failed to locate any CGI in the promoter region (Figure S3A). However, we found that one CGI was in the intron 2 (Intron-CGI); and the remaining one CGI overlapped with exon 4 (Ex-CGI). Studies suggested that one of the roles of gene body methylation is the regulation of transcription process (Aran et al., 2011; Maunakea et al., 2010; Singer et al., 2015). Hence, we hypothesized that Ex-CGI methylation on HOTAIR gene promoted the transcription process of HOTAIR RNA. To show that Ex-CGI hypermethylation was associated with high HOTAIR levels, quantitative methylation specific PCR (qMSP) was conducted to show that Ex-CGI was frequently methylated in a majority of PDAC cell lines, but not in the non-tumor HPDE cells and PDAC CFPAC-1 cells (Figure 1B). It was consistent with HOTAIR expression in PDAC cells in which both HPDE and CFPAC-1 cells had low HOTAIR level. In addition, most HCC, CRC and breast cancer cells had high expression of HOTAIR, while their Ex-CGI were hypermethylated (Figure 1C-E, Figure S1E, F). In contrast, low Ex-CGI methylation level was observed in MIHA, DLD-1 and MDA-MB-415 cells, which expressed relatively low HOTAIR level.

**Fig. 1.**
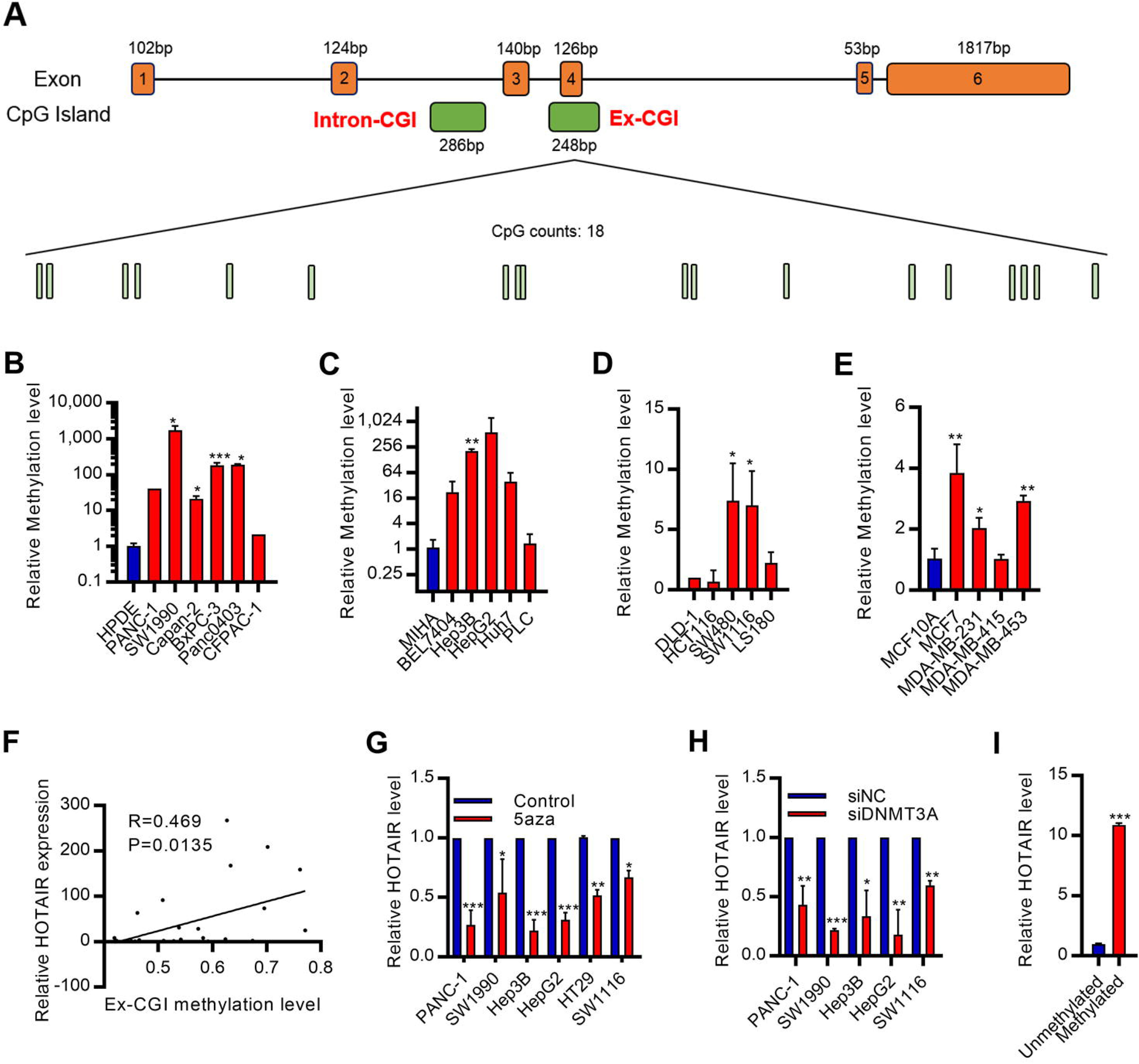
Methylation of Ex-CGI was associated with HOTAIR expression in cancer. **(A)** Schematic of CpG island (CGI) distribution in HOTAIR gene. Two CGI was found in the intragenic region of HOTAIR. One CGI was in the intron 2 (Intron-CGI); and the remaining one CGI overlapped with exon 4 (Ex-CGI). 18 CpG sites were found in the Ex-CGI. **(B-E)** qMSP analysis of methylation level of Ex-CGI in **(B)** PDAC, **(C)** HCC, **(D)** CRC and **(E)** breast cancer cells. **(F)** Pyrosequencing analysis of the correlation between Ex-CGI methylation level and HOTAIR expression in PDAC tumors. **(G-H)** Changes in HOTAIR expression level after **(G)** treatment with 5aza or **(H)** knockdown of DNMT3A in PDAC, HCC and CRC cells. **(I)** *In vitro* methylation assay for analysis of HOTAIR expression after transfecting pEGFP-N1 with methylated HOTAIR gene in Cos7 cells where endogenous human HOTAIR was not expressed. Data are from at least three independent experiments and plotted as means ± SD. * *P* < 0.05, ** *P* < 0.01, *** *P* < 0.001.

To investigate the clinical importance of Ex-CGI in PDAC primary tumors, qMSP at the HOTAIR Ex-CGI was performed in human PDAC primary tumors. We found that Ex-CGI of human PDAC primary tumors were significantly hypermethylated (Figure S3B). Subsequently, pyrosequencing had confirmed that several CpG sites at Ex-CGI were frequently hypermethylated and were positively associated with HOTAIR expression level in PDAC primary tumors (Figure 1F, Figure S3C). Analysis of The Cancer Genome Atlas (TCGA) database also revealed the upregulation of Ex-CGI methylation level and its positive correlation with HOTAIR expression in cancer (Figure S4). We next investigated the effect of Ex-CGI methylation on HOTAIR expression. We found that DNA demethylation using 5-aza-2’-deoxycytidine (5-aza) on PDAC, HCC and CRC cells resulted in a decrease in HOTAIR expression (Figure 1G, Figure S5A). Conversely, *in vitro* methylation of HOTAIR gene by SssI DNA methyltransferase induced the expression of HOTAIR in COS7 cells where endogenous human HOTAIR was not expressed (Figure 1I). Collectively, our results suggested that *de novo* DNA methylation in Ex-CGI contributed to HOTAIR expression in cancer.

### DNA methylation in Exon-CpG Island promoted transcription elongation of HOTAIR gene

Previous studies suggested that intragenic DNA methylation altered chromatin structure and transcription process (Lorincz et al., 2004). Therefore, we hypothesized that Ex-CGI enhances HOTAIR transcription through stabilizing protein factors on the intragenic regions. We found that although transcription of Exon 2 occurred in both non-tumor and cancer cells, transcription after Ex-CGI was greatly hindered in HPDE and CFPAC-1 cells (Figure 2A, B). ChIP assay also revealed that Ser2-phosphorylated RNA PolII, which was associated with transcription elongation (Komarnitsky et al., 2000), was significantly enriched in Ex-CGI in PDAC, HCC and CRC cells with high Ex-CGI methylation level and HOTAIR expression (Figure 2C). The association between Ex-CGI methylation and chromatin accessibility was also observed in PDAC cells (Figure 2D). In contrast, DNA demethylation in Ex-CGI significantly reduced the occupancy of Ser2-phosphorylated RNA PolII, chromatin accessibility and transcription efficiency downstream Ex-CGI (Figure 2E-G). This suggested that Ex-CGI methylation played a role in stabilizing HOTAIR transcription process. In non-tumor cells or cells with unmethylated Ex-CGI, transcription of HOTAIR will be terminated when RNA PolII read to the exon harbouring Ex-CGI, which allowed the complete transcription of HOTAIR RNA.

**Fig. 2.**
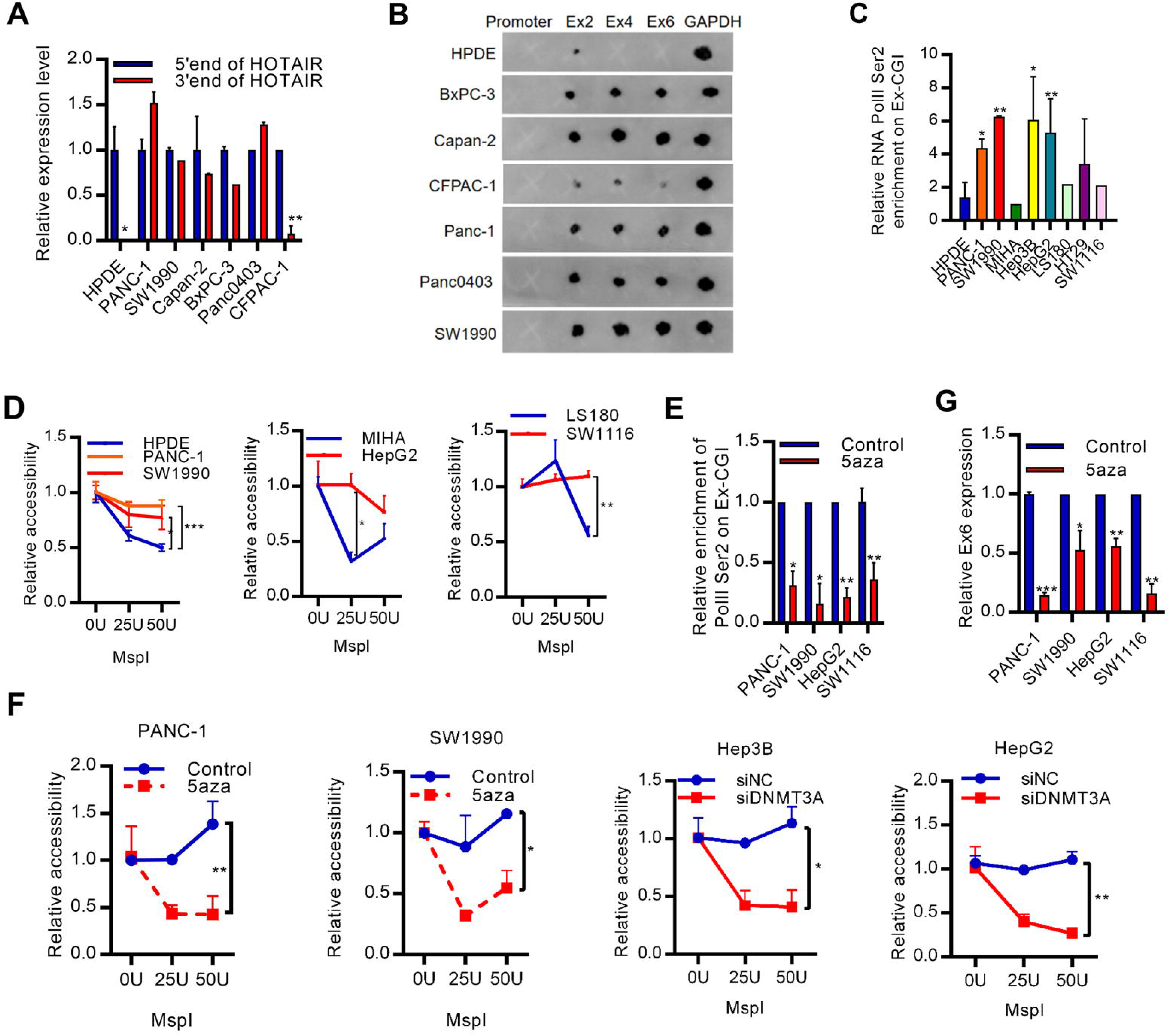
Methylation of Ex-CGI promotes HOTAIR expression through facilitating its transcription elongation. **(A)** Expression pattern of 5’ end and 3’ end of HOTAIR in PDAC cell lines. **(B)** Transcription elongation efficiency of HOTAIR in PDAC cells by nuclear run on assay, followed by northern dot plot. **(C)** ChIP analysis for RNA PolII Ser2 on Ex-CGI in PDAC, HCC and CRC cells. **(D)** Chromatin structure along HOTAIR gene in PDAC, HCC and CRC cells by MspI chromatin accessibility assay. MspI mimics the binding of RNA PolII to HOTAIR gene. Compared to promoter, the accessibility to exon 6 was reduced, was similar in cancer cells with high HOTAIR expression. **(E)** ChIP analysis for RNA PolII Ser2 on HOTAIR gene after 5aza treatment in PANC-1 and SW1990 cells. **(F)** Chromatin structure along HOTAIR gene after inhibition of DNMT in PDAC, HCC and CRC cells. **(G)** Transcription elongation efficiency of HOTAIR after inhibition of DNMT in PDAC and HCC cells. Data are from at least three independent experiments and plotted as means ± SD. * P < 0.05, * * P < 0.01, * * * P < 0.001.

### MLL1-mediated H3K4me3 promoted the establishment of Exon-CpG Island methylation

*De novo* exonic DNA methylation pattern is established by DNA methyltransferase DNMT3A and DNMT3B (Okano et al., 1999). DNMT3B recognized intragenic histone modifications, which were the marks of active transcription elongation process (Morselli et al., 2015; Wen et al., 2014), to promote *de novo* gene body methylation (Dhayalan et al., 2010; Hahn et al., 2011). To investigate histone modification-mediated Ex-CGI methylation, we performed ChIP assay to profile the histone modification status along the HOTAIR gene (Figure 3A, B). We found that H3K4me3 was enriched in Ex-CGI. Also, knockdown of H3K4 methyltransferase MLL1 inhibited H3K4me3 occupancy, Ex-CGI methylation, occupancy of Ser2-phosphorylated RNA PolII and HOTAIR expression in PDAC, HCC and CRC cells (Figure 3C-E). On the other hand, upregulation of DNMT3A occupancy, but not DNMT3B nor DNMT1, was observed on Ex-CGI in cancer cells (Figure S5B-D). Consistently, only knockdown of DNMT3A inhibited HOTAIR transcription (Figure 1H, Figure S5E-G). Collectively, our results suggested that H3K4me3 promoted DNMT3A-mediated Ex-CGI methylation and HOTAIR expression.

**Fig. 3.**
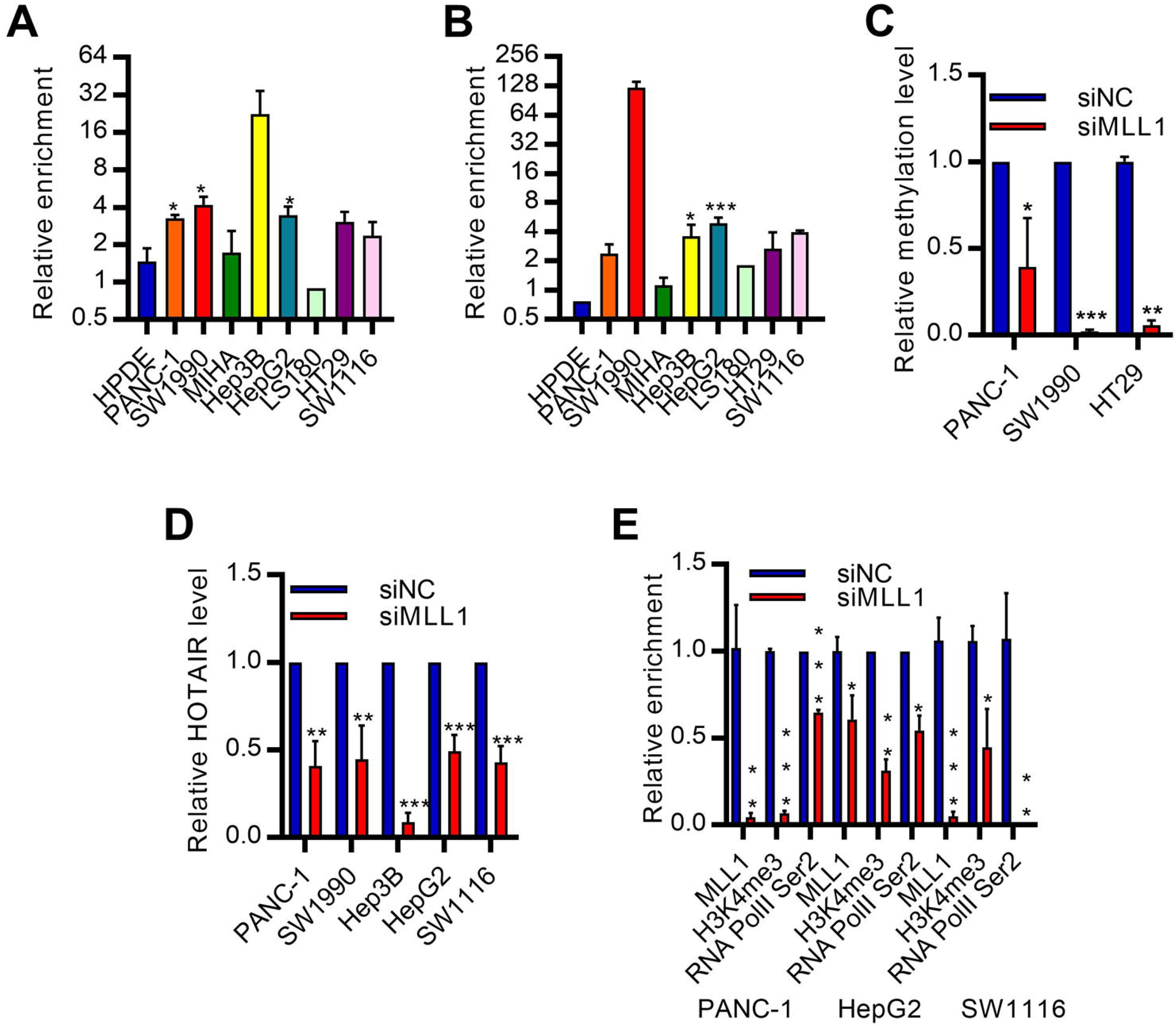
MLL1-H3K4me3 promotes Ex-CGI methylation and HOTAIR expression. **(A, B)** ChIP analysis revealed the occupancies of (**A**) H3K4me3 and (**B**) MLL1 on Ex-CGI in PDAC, HCC and CRC cells. Enrichment level of H3K4me3 and MLL1 in cancer cells were compared to corresponding non-tumor cells. **(C, D)** (**C**) Methylation level of Ex-CGI and (**D**) HOTAIR expression after knockdown of MLL1 in PDAC, HCC and CRC cells. **(E)** MLL1, H3K4me3 and RNA PolII Ser2 occupancies on HOTAIR Ex-CGI after knockdown of MLL1. Data are from at least three independent experiments and plotted as means ± SD. * P < 0.05, * * P < 0.01, * * * P < 0.001

### CDK7-CDK9 axis promoted HOTAIR expression by induction of H3K4 methylation and Ex-CGI methylation

We further explored the regulatory mechanism of MLL1-H3K4me3-Ex-CGI methylation. Preceding works suggested the role of positive transcription elongation factor b (p-TEFb) on promoting transcription elongation through indirectly mediating epigenetic changes in coding region (Eissenberg et al., 2006; Tanny, 2014). We hypothesized that p-TEFb promoted the methylation of H3K4me3-Ex-CGI to facilitate transcription of HOTAIR in PDAC, HCC and CRC cells. First, pharmacological suppression of p-TEFb by DRB inhibited HOTAIR expression (Figure S6A). Also, inhibition of p-TEFb regulator Brd4 (Itzen et al., 2014), which promoted PDAC progression (Wang et al., 2015), by PFI-2 inhibited HOTAIR expression (Figure S6B). SiRNA-mediated inhibition and pharmacological inhibition of CDK9, which is the key component of p-TEFb, by LDC-067 (Albert et al., 2014) in PDAC, HCC and CRC resulted in a reduction of Ex-CGI methylation and HOTAIR expression (Figure 4A, Figure S6D-E). The lack of CDK9 activity also impaired the transcription elongation process, resulting in the failure in producing full length functional HOTAIR transcript (Figure 4B-E). Since CDK9 can be regulated by multiple pathways, which target different phosphorylation sites on CDK9 (Nekhai et al., 2014), we next investigated which CDK9 regulator was involved in HOTAIR expression. Knockdown of CDK7 (but not other CDK9 regulator) inhibited HOTAIR expression (Figure S7A-D). CDK7 is frequently upregulated and promotes carcinogenesis through mediating transcriptional activation of oncogenes (Greenall et al., 2017; Kalan et al., 2017; Kwiatkowski et al., 2014; Wang et al., 2015). As expected, pharmacological inhibition of CDK7 by THZ-1 (Albert et al., 2014) in PDAC, HCC and CRC cells inhibited Ex-CGI methylation and HOTAIR expression (Figure 5A, Figure S7E).

**Fig. 4.**
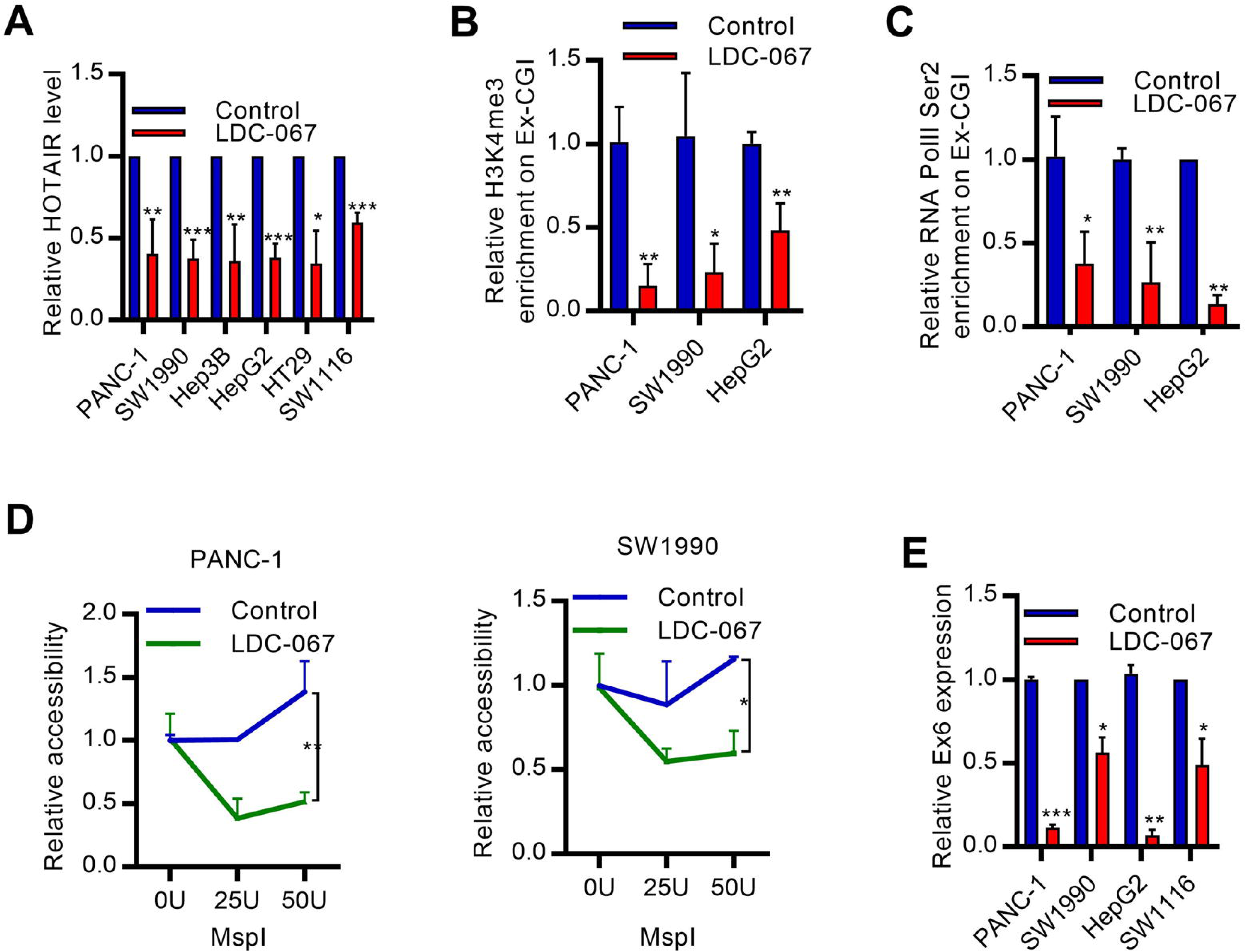
CDK9 promotes transcription of HOTAIR in cancers. **(A)** HOTAIR expression after inhibiting CDK9 by LDC-067 in PDAC, HCC and CRC cells. **(B-C)** ChIP analysis revealed the occupancies of **(B)** RNA PolII Ser2 and **(C)** H3K4me3 and on Ex-CGI after inhibiting CDK9. **(D)** Chromatin structure along HOTAIR gene after inhibition of CDK9 in PDAC, HCC and CRC cells. MspI mimics the binding of RNA PolII to HOTAIR gene. The accessibility to exon 6 was reduced, compared to promoter. **(E)** Nuclear run-on qPCR analysis revealed transcription elongation efficiency of HOTAIR after inhibition of CDK9 in PDAC and HCC cells. The expression of Exon 6 was compared to expression of Exon 2. Transcription efficiency of Exon 6 was decreased after inhibition of CDK9. Data are from at least three independent experiments and plotted as means ± SD. * P < 0.05, * * P < 0.01, * * * P < 0.001

**Fig. 5.**
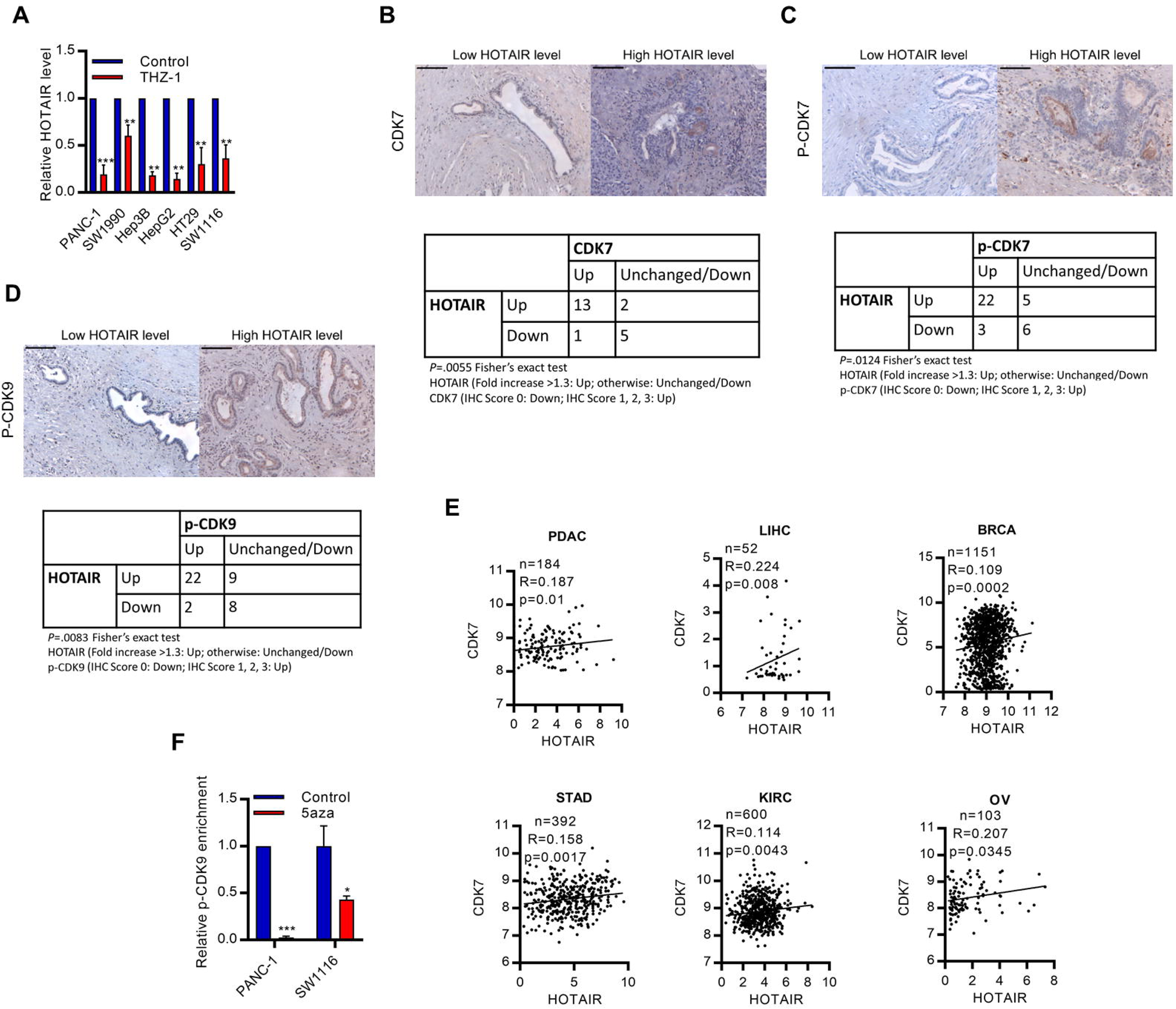
The upregulated of CDK7-CDK9 axis promotes HOTAIR expression. **(A)** HOTAIR expression after inhibiting CDK7 by THZ-1 in PDAC, HCC and CRC cells. **(B-D**) IHC for (**B**) CDK7 expression, (**C**) CDK7 (T170) phosphorylation and (**D**) CDK9 (T170) phosphorylation in PDAC tumor sections with (left) low and (middle) high HOTAIR expression level. (right) Protein expression in PDAC primary tumors was compared with adjacent non-tumor tissues and is correlated with HOTAIR expression. **(E)** The correlation between CDK7 expression and HOTAIR expression in publicly available expression dataset (TCGA) of PDAC, LIHC and BRCA; n = 1,380 samples. **(F)** ChIP analysis for CDK9 (T170) phosphorylation on Ex-CGI in PDAC and CRC cells after 5aza treatment. Data are from at least three independent experiments and plotted as means ± SD. * P < 0.05, * * P < 0.01, * * * P < 0.001.

Furthermore, to study the clinical significance of CDK7-CDK9-HOTAIR axis, we examined the phosphorylation level of CDK7 and CDK9 in PDAC cells and primary tumors. We found that CDK7 and CDK9 were robustly phosphorylated in PDAC cells and primary tumors and were associated with HOTAIR expression level (Figure 5B-D, Figure S7F). Analysis of TCGA database also revealed the correlation between CDK7 and HOTAIR expression level in cancer (Figure 5E). More importantly, we demonstrated that DNA methylation played a role in recruiting CDK9 to the HOTAIR gene. DNA demethylation induced by 5-azacytidine could reduce the enrichment levels of p-CDK9 at the gene body of HOTAIR (Figure 5F). As such, it suggested that gene body methylation at the exon of HOTAIR could promoted the binding of CDK9, and enhanced the elongation process via RNA PolII phosphorylation. The positive feedback loop involving the methylation of H3K4 and exon CpG island and the recruitment of CDK9 would then be established, which allowed the persistent expression of HOTAIR. Collectively, these results demonstrated CDK7-CDK9 axis contributed to the establishment of positive feedback loop that corporate phosphorylation of RNA PolII, induction of H3K4 methylation, Ex-CGI methylation and recruitment of more CDK9, and in turn promoted persistent HOTAIR overexpression (Figure 6).

**Fig. 6.**
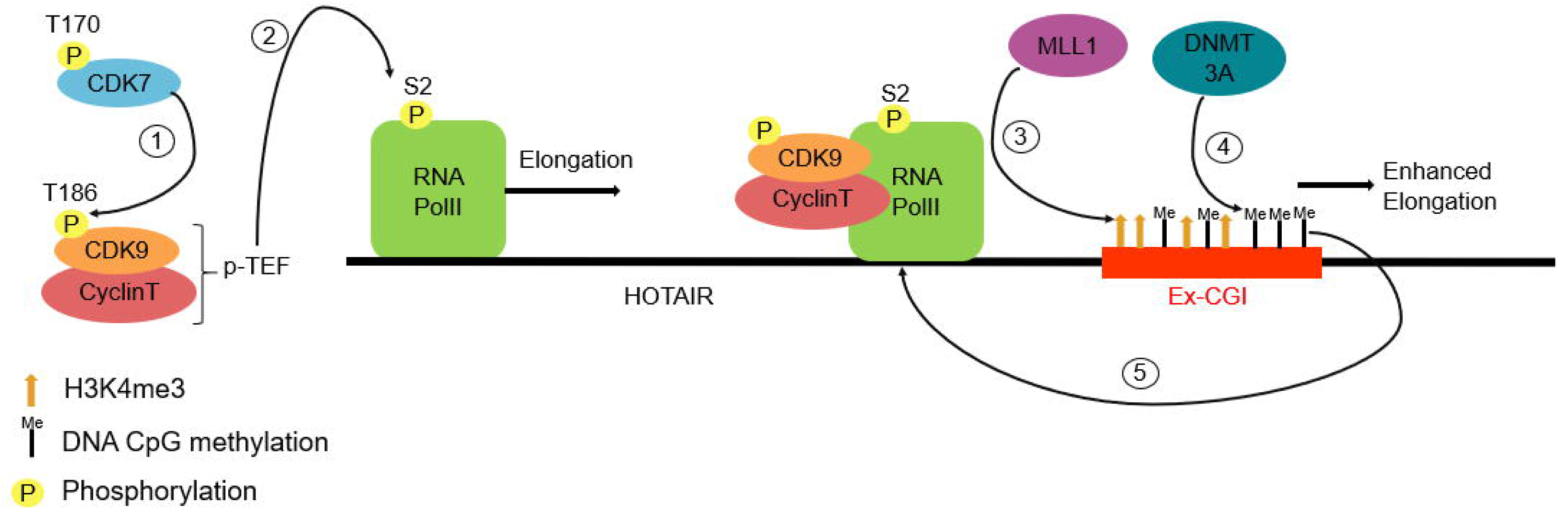
Schematic diagram describing the role of CDK7-CDK9 mediated intragenic H3K4me3 and DNA methylation in promoting HOTAIR transcription in cancer. Phosphorylation of oncogenic CDK7 (T170) activates CDK9, which is the key component of p-TEFb, by phosphorylating T186, which in turn promoted the elongation process of HOTAIR by activating RNA Polymerase II Ser2. The Ex-CGI methylation in the HOTAIR gene was regulated by MLL1-mediated H3K4 methylation during transcription elongation process. The methylation of Ex-CGI enhanced the elongation process, resulting in upregulation of HOTAIR in cancer.

## DISCUSSION

HOTAIR, which is a well-characterized lncRNA, is upregulated in many cancer types. Studies have demonstrated that HOTAIR played important in promoting cancer progression by regulating gene expression (Tang et al., 2018). HOTAIR guilds EZH2/PRC2 complex to tumor suppressor for triggering gene silencing H3K27me3 (Li et al., 2017). Also, HOTAIR interacts with H3K4 demethylase LSD1 and induces gene silencing (Zhang et al., 2015). However, the mechanism contributing to the deregulation of HOTAIR in cancers is still largely unknown. Here, we found that methylation of Ex-CGI, which is in exon 4 of HOTAIR gene, promoted HOTAIR expression by facilitating the transcription of full-length transcript. Important, we demonstrated the intragenic DNA methylation was regulated by CDK7-CDK9-RNA PolII-mediated H3K4me3.

DNA methylation at the cytosine residues has been reported to be an important regulator of gene expression. Methylation at the promoter frequently links to gene silencing. However, intragenic or specifically exon methylation may result in either gene activation or gene silencing. DNA methylation at the first exon inhibits gene expression by blocking transcription initiation (Brenet et al., 2011; Jjingo et al., 2012). Study found that intragenic regions with DNA methylation were depleted by RNA PolII and histone markers of active transcription elongation (Lorincz et al., 2004). However, another study suggested that intragenic DNA methylation facilitated gene elongation process (Jjingo et al., 2012). Here, we observed the positive correlation between Ex-CGI and HOTAIR expression in many cancer types. In addition, depletion of Ex-CGI methylation resulted in the formation of heterochromatin, and in turn impaired RNA PolII transcription elongation process and decreased in HOTAIR expression, while *in vitro* methylation of Ex-CGI increased HOTAIR expression. These results might suggest that gene body methylation promoted HOTAIR expression in cancer by facilitating transcription elongation process.

Although many studies suggested that intragenic DNA methylation was involved in gene regulation, how it is regulated is largely unexplored. DNA methylation is frequently linked to histone modifications in many cancers (Kondo, 2009). For instance, DNA methylation and H3K9 methylation are often co-exist on same promoter, resulting in gene silencing (Baylin et al., 2016). A previous report also hypothesized that DNMT recognized histone modification to create *de novo* DNA methylation (Tanny, 2014). Studies found that DNMT3A and DNMT3B recognized H3K36me3, which is a marks of active transcription elongation process, to promote *de novo* intragenic methylation (Dhayalan et al., 2010; Singer et al., 2015). Here, we demonstrated DNMT3A, but not DNMT3B, recognized the H3K4me3, which was also a hallmark histone modification of active transcription elongation (Chen et al., 2015), to promote Ex-CGI methylation and HOTAIR expression.

CDK9, a key component of p-TEFb, and its regulator CDK7 are fundamentally important for formation of transcription elongation complex, releasing the RNA PolII from the promoter and triggering transcription elongation processes. CDK9 phosphorylates the Ser2 position of Rpb1 CTD, which is the regulatory unit of RNA PolII, for the recruiting of other members of the transcription elongation complex, including WAC and PAF (Tanny, 2014). These also contribute to the recruitment of histone modification enzymes such as SETD1 for intragenic H3K36 methylation (Eissenberg et al., 2007). Also, a recent finding also demonstrated the link between CDK9 and intragenic histone modification markers. The oncogenic CDK9 phosphorylated BRG1, resulting in establishment of gene silencing H3K9me2 (Zhang et al., 2018). In this study, we observed the positive correlation between CDK7, CDK9 and HOTAIR level in cancers. Also, inhibition of these transcription elongation regulator inhibited Ex-CGI methylation and HOTAIR expression. Importantly, depletion of Ex-CGI methylation reduced the enrichment levels of p-CDK9 at the gene body of HOTAIR. These might suggest the presence of feedback loop that intragenic DNA methylation recruited CDK9 for the persistent expression of HOTAIR.

In summary, we unraveled a novel molecular pathway in upregulating oncogenic lncRNA in cancers, which linked up gene body histone methylation, DNA methylation and RNA transcription elongation. We revealed for the first time that CDK7-CDK9-H3K4me3 axis regulated Ex-CGI methylation, and then led to a feedback loop to promote the recruitment of CDK9 to HOTAIR gene, and subsequently promoted HOTAIR expression in cancer. The identification of this pathway could provide the explanation to how certain oncogenic lncRNAs were ubiquitously overexpressed in multiple cancer types that showed diverse genetic profiles.

## ACKNOWLEDGMENTS

The research was supported by General Research Fund, Research Grants Council of Hong Kong [CUHK462713, 14102714, 4171217 and 14120618 to Y.C.]; National Natural Science Foundation of China [81672323 to Y.C.]; Direct Grant from CUHK to YC. Microarray data are available at ArrayExpress E-MTAB-7305.

## AUTHOR CONTRIBUTIONS

C.H.W. and C.H.L. designed and performed the experiments and drafted the manuscript. Q.H. assisted to obtain data. J.H.M.T. and K.F.T. provided some research materials. Y.C. was the PI of the grant, overlook the whole progress and revised the manuscript.

## DECLARATION OF INTEREST

The authors declare no competing interests.

## STAR METHODS

### METHOD DETAILS

#### Cell Culture and Drug treatment

PDAC Cell lines PANC-1, SW1990, CAPAN-2, CFPAC-1, PANC0403, and BxPC-3; HCC cell lines HepG2, Hep3B, and PLC/PRF/5 (PLC); CRC cell lines DLD-1, SW1116, SW480, HCT116, HT29 and LS180; and HEK293T were purchased from American Type Culture Collection. Human pancreatic ductal epithelial (HPDE) cell line was a gifted from from Dr Ming-Sound Tsao (University Health Network, Ontario Cancer Institute and Princess Margaret Hospital Site, Toronto). The non-tumorigenic human hepatocyte cell line MIHA was kindly provided by Dr. J.R. Chowdhury’s laboratory at Albert Einstein College of Medicine. The human HCC cell line Huh7 was obtained from Dr. H. Nakabayashi, Hokkaido University School of Medicine, Sapporo, Japan and Bel-7404 was obtained from Cell Bank of the Chinese Academy of Science. HPDE cells were cultured in Keratinocyte Serum free medium supplemented with 50 µg/mL bovine pituitary extract, 0.2 ng/mL human epithelial growth factor (Invitrogen) and 3% antibiotic and antimycotic (Gibco). BxPC-3 cells were maintained in RPMI-1640 supplemented with 10% FBS and 100 units/ml penicillin and 100 µg/ml streptomycin (Gibco). The remaining cell lines were maintained in DMEM containing 10% FBS, 100 units/ml penicillin and 100 µg/ml streptomycin. For drug treatment, cells were treated with 5-Aza-2′-deoxycytidine (5aza) (Sigma), THZ1 (MedChem Express), LDC-067 (MedChem Express), PFI-2 (ApexBio), DRB (ApexBio) for 72 h before the experiment as described.

#### Clinical Sample and Histology

Sixty pairs of PDAC tumor and adjacent non-tumor tissues were obtained from patients who underwent pancreatic resection at the Prince of Wales Hospital. All specimens were fixed and embedded into paraffin. PDAC tissue specimens were fixed in 4% buffered formalin for 24 h and stored in 70% ethanol until paraffin embedding. 5-μm sections were stained with haematoxylin and eosin (HE) to locate tumors and adjacent normal tissues.

#### Microarray analysis

Microarray analysis of lncRNA expression on 4 pairs of PDAC tumor samples was performed on Arraystar Human LncRNA Microarray V4.0 platform. Microarray data are available at ArrayExpress E-MTAB-7305.

#### Constructs and Cell transduction

Lentiviral vector with full length of HOTAIR was synthesized by abm. To achieve stable ectopic expression, lentivirus was produced by cotransfecting HEK293T cells with HOTAIR-containing lentiviral vector and 3 packaging vectors: pMDLg/pRRE, pRSV-REV, and pCMV-VSVG as previously described (Chen et al., 2007). Then HPDE cells were transduced with by lentivirus with polybrene for 6 h, followed by antibiotic selection. The efficiency of HOTAIR overexpression was validated by qRT-PCR.

#### siRNA transfection

siRNAs targeting HOTAIR, CDK2, CDK7, CDK9, PP1, DNMT1, DNMT3A, DNMT3B and MLL1 were purchased from GenePharma and were dissolved in siRNA buffer (Thermo Fisher). For transient knockdown, cells were transfected with siRNAs using Lipofectamine 3000 transfection reagent (Thermo Fisher) for 72 h. Cells were then collected for the experiments as described.

#### Luciferase assay

Luciferase report plasmid was constructed by cloning 500 bp, 1 kb and 2 kb upstream of HOTAIR transcription start site into pGL3-basic plasmid. HEK293T cells were transfected with the luciferase reporter plasmid, control vector (pGL3-Control) and Renilla luciferase reporter plasmid using Lipofectamine 3000 transfection reagent and P3000 Reagent (Invitrogen). Luciferase activity was analyzed by Dual Luciferase Reporter Assay kit (Promega).

#### Quantitative reverse transcription PCR (qRT-PCR)

Cell lines total RNA was extracted by TRIzol Reagent (Invitrogen). Formalin-fixed, paraffin-embedded (FFPE) sample RNA was isolated by miRNeasy FFPE Kit (Qiagen) according to manufacturer’s protocol. Measurement of gene expression level was performed by qRTPCR. cDNA was synthesized using High-Capacity cDNA Reverse Transcription Kit (Applied Biosystems) according to manufacturer’s instructions. Quantitative PCR was performed by ABI 7900HT RealTime PCR system (Applied Biosystems) using SYBR Green PCR Master Mix (Applied Biosystems). The PCR primer sequences are listed in Table S1.

#### MTT cell viability assay

A 3-(4, 5-dimethylthiazol-2-yl)-2, 5-diphenyltetrazolium bromide MTT assay was performed measured cell viability. To investigate the effect of HOTAIR on carcinogenesis, cells were transfected with siRNA targeting HOTAIR for 72 h. To determine effective concentration of each drug, cells were incubated with a series of dosage of drug for 72 h. Then, cells were incubated with 0.65 mg/mL MTT at 37 °C for 2 h. DMSO was used to dissolve formazon crystals and absorbance was measured at 595 nm.

#### Colony formation assay

Anchorage-independent colony formation assay was performed as described previously (Li et al., 2015) to study the transformation ability of HPDE when HOTAIR is ectopically expressed

#### Chromatin immunoprecipitation (ChIP)

ChIP was performed using EZ-Magna ChIP HiSens kit (Millipore) according to manufacturer’s protocol. Briefly, cells were cross-linked with 1% formaldehyde for 10 min and the reaction was stopped with 125 mM glycine. Chromatin was then isolated and sonicated into fragments of 200-600 bp. Cross-linked chromatin was incubated at 4°C overnight anti-IgG (-ve control), anti-RNA Polymerase II Phosphorylated Ser5 (Millipore, 05-623), anti-RNA Polymerase II Phosphorylated Ser2 (Covance, MMS-129R), anti-MLL1 antibody (Millipore, ABE240), anti-H3K4me3 antibody (Millipore, CS200580), anti-DNMT1 antibody (Cell Signaling, 5119), anti-DNMT3A antibody (Cell Signaling, 2160), and anti-DNMT3B antibody (Cell Signaling, 2161). The precipitated DNA was quantitated by qPCR.

#### quantitative Methylation-Specific PCR (qMSP)

Cell lines DNA was isolated by NK Lysis Buffer (Chen et al., 2015). FFPE sample DNA was isolated by QIAamp DNA FFPE Tissue Kit (Qiagen) according to manufacturer’s protocol. Bisulphite conversion was performed using the EZ DNA Methylation-Lightning™ Kit (Zymo Research). 500ng Genomic DNA was added to cytosine-to-uracil conversion reagent and was placed in a thermal cycler at 98 °C for 8 min and 54°C for 60 min, followed by in-column desulphonation at room temperature for 20 min. The methylation state of HOTTIP CpG island was analyzed by both PCR and qPCR using methylation specific primers and unmethylation specific primers.

#### Pyrosequencing

Each pyrosequecing primer set consisted of one unlabeled forward primer, one biotinylated reverse primer and one sequencing primer. Four sequencing primers specific to bisulphite-treated Exon CpG island were used to analyze methylation status of each CpG site. Pyrosequencing was performed on the PSQ 96MA system (Qiagen) and the resulting methylation percentage of cytosine in each CpG dinucleotide was calculated by Pyro Q-CpG™ software.

#### *In vitro* DNA methylation assay

HOTAIR gene was cloned into expression vector pEGFP-N1. In vitro DNA methylation was performed by CpG Methyltransferase (M.SssI) (NEB) in the presence of 320 μM S-adenosylmethionine (NEB). pEGFP-N1 with HOTAIR being methylated was transfected in Cos7 cells using Lipofectamine 3000 Reagent and P3000 Reagent. RNA was extracted by TRIZOL and analyzed by qRT-PCR.

#### Nuclear Run-on assay

Nuclear run-on assay was performed according to reported protocols with modifications (Roberts et al., 2015). Cells were washed twice with ice-cold PBS, scraped and pelleted by centrifugation at 800g 4°C for 10 mins. Nuclei was isolated by NP-40 lysis buffer with proteinase inhibitors cocktail (Roche) and phosphatase inhibitor cocktail (Thermo Fisher), followed by centrifugation at 800g 4°C for 5 mins, and was resuspended in 120 μL nuclei storage buffer (40% glycerol, 5mM MgCl2, 0.1mM EDTA and 50mM Tris, pH 8.0). Nuclear Run-on assay was performed by incubating the nuclei at 37 °C for 30 mins in transcription buffer (2.5 U RNase inhibitor (Applied Biosystem), 7 % glycerol 5mM MgCl_2_, 75mM KCl, 50μM EDTA, 25mM Tris and 25mM HEPES, pH 7.5) containing nucleoside triphosphates (NTPs, 0.35 mM ATP, GTP and CTP, 0.4 μM UTP) and 0.25 mM DIG-labelled uridine 5’-triphosphate (UTP) (Roche), followed by RNA isolation by TRIZOL reagent. Nascent transcripted RNA was analyzed by either dot-blot or qPCR. For dot-blot analysis, RNA probe specific to HOTAIR exons and GAPDH were blotted on the positive charged nylon membrane (Roche) for 1 h. The blot was washed with SSC buffer (0.3 M NaCl, 30mM sodium citrate, pH7), followed by incubation at 80°C for 2 h. Then, the blot was incubated with hybridization buffer (50% formamide, 10X Denhardt’s solution, 0.2M NaCl, 1mM EDTA, 20mM Tris, pH8.0) at 65°C for 1 h. Before hybridization, DIG-labelled RNA was heated at 95°C for 10 mins to degrade RNA into small fragments. RNA sample was hybridized to RNA probe overnight. After hybridization, the blot was washed three times with SSC buffer and was incubated with anti-DIG antibody (abcam, ab119345) for 1 h at room temperature. The blot was washed with SSC buffer and the hybridized RNA was visualized by Novex AP Chemiluminescent Substrate (CDP-Star) (Thermo Fisher). For qPCR analysis, DIG-labelled RNA was immunoprecipitated with anti-DIG antibody (abcam ab119345) labelled protein A/G magnetic beads according to previous report. Briefly, the protein A/G magnetic beads were conjugated with anti-DIG antibody for 10 mins at room temperature, followed by blocking with Denhardt’s solution. DIG-labelled RNA was immunoprecipitated for 30 mins at room temperature. RNA was extracted by TRIZOL and analyzed by qRT-PCR as described.

#### Chromatin Accessibility assay

The accessibility of HOTAIR gene was assayed with MspI digestion, which mimicked the binding of RNA Polymerase II, according to previous report with minor modifications (Nguyen et al., 2001). Briefly, 10^7^ cells were washed with cold PBS twice. The cell pellet was resuspended in nuclei isolation buffer (10mM NaCl, 3mM MgCl2, 1mM PMSF, 1% NP-40, and 10mM Tris, pH 7.4) and was incubated on ice for 10 minutes. The cells were homogenized on ice by Dounce homogenizer. The nuclei were pellet by centrifugation at 1000 g, 10 mins, 4°C and resuspended in MspI buffer. 0U, 25U and 50U MspI (NEB) was added to digest 1 × 10^6^ nuclei at 37°C for 1h. DNA was extracted by phenol: chloroform: isoamylalcohol, followed by ethanol participation in the presence of sodium acetate. The relative accessibility of 3’ end to 5’ end of HOTAIR gene was analyzed by qPCR.

#### Immunoblot analysis

Whole cell extract was prepared by lysing cells in NP-40 lysis buffer with proteinase inhibitors and phosphatase inhibitor. Protein concentration was determined by BCA assay (Pierce). Proteins were resolved by SDS-PAGE at different percentages, transferred to PVDF membrane and immunoblotted overnight at 4 °C with antibodies against CDK7 (phosphor T170) (rabbit; abcam ab155976; 1:1000), CDK9 (phosphor T186) (rabbit; Cell signaling 2459; 1:1000) and GAPDH (rabbit; Cell signaling 5174; 1:1000). Then, the blots were washed three times with TBST, followed by incubation with 1:2000 secondary anti-rabbit antibody at room temperature for 1 h. Images of immunoblot were taken with ChemiDoc Imaging System (Bio-Rad).

#### Immunohistochemistry

Immunostaining was performed using 5-μm PDAC FFPE tissue sections. Antigen retrieval was carried out using PT module (Thermo Fisher). The sectioned tissues were deparaffinized and rehydrated by xylene and a series of graded ethanol. Antigen retrieval was carried out using PT module (Thermo Fisher, Waltham, MA). Then, immunostaining was performed using Histostain-Plus IHC Kit, HRP, broad spectrum (Life Technologies, Carlsbad, CA) according to manufacturer’s protocol. Antibodies against CDK7 (rabbit; abcam ab216437; 1:100), CDK7 (phospho T170) (rabbit; abcam ab59987; 1:50), CDK9 (phospho T186) (rabbit; Cell signaling 2459; 1:100) were used for staining at 4°C overnight. Sections were counter-stained with hematoxylin. Images of immunohistochemistry were taken with Spot Digital Camera & Leica Microscope Biological Imaging System (10X and 40X magnification). A scoring system, based on the percentage of positive cells and staining intensity under 10X magnification, was used to quantify the staining. 4 categories (0, 1, 2, and 3) were demoted as 0%, 1-10%, 10-50%, and >50%. The staining intensity of the tumor section was compared to adjacent non-tumor tissue. The association between HOTAIR expression and intensity of CDK7, CDK7 (phosphor T170) and CDK9 (phosphor T186) protein staining was performed using two-tailed Fisher’s exact t-test.

#### Analysis of patient data sets

Publicly available patient datasets (Pancreatic Ductal Adenocarcinoma, liver hepatocellular carcinoma, colorectal adenocarcinoma, breast carcinoma, lung adenocarcinoma, head and neck squamous cell carcinoma) was obtained from The Cancer Genome Atlas (TCGA). Processed data in Z-score were used in analysis of CDK7 and HOTAIR expression. Processed data in beta value were used in analysis of Ex-CGI methylation level. For the association analysis between HOTAIR expression and Ex-CGI methylation level; and between HOTAIR expression and CDK7 expression, linear regression was used.

#### Statistical analysis

GraphPad Prism 6 (GraphPad Software) was used for statistical analysis. Unpaired two-tailed student’s t-test was used to compare the differences between two groups, unless noted otherwise. Data are represented as mean ± s.d. *P* value less than 0.05 was considered statistically significant. For association study between CGI methylation and HOTAIR expression level in PDAC tissues, we first calculated the beta value of each CpG site in each sample (Du et al., 2001) and compared with the beta value of their corresponding adjacent non-tumor tissue. Then we use the mean beta value in each pair of PDAC tissue for correlation analysis with their corresponding relative HOTAIR expression.

## SUPPLEMENTAL INFORMATION

Supplementary Information including Methods and Materials and nine Supplementary Figures.

