## Supplementary material for "The Establishment of CDK9/ RNA PolII/H3K4me3/DNA Methylation Feedback Promotes HOTAIR Expression by RNA Elongation Enhancement in Cancer": suppl data

**Supplementary Information**

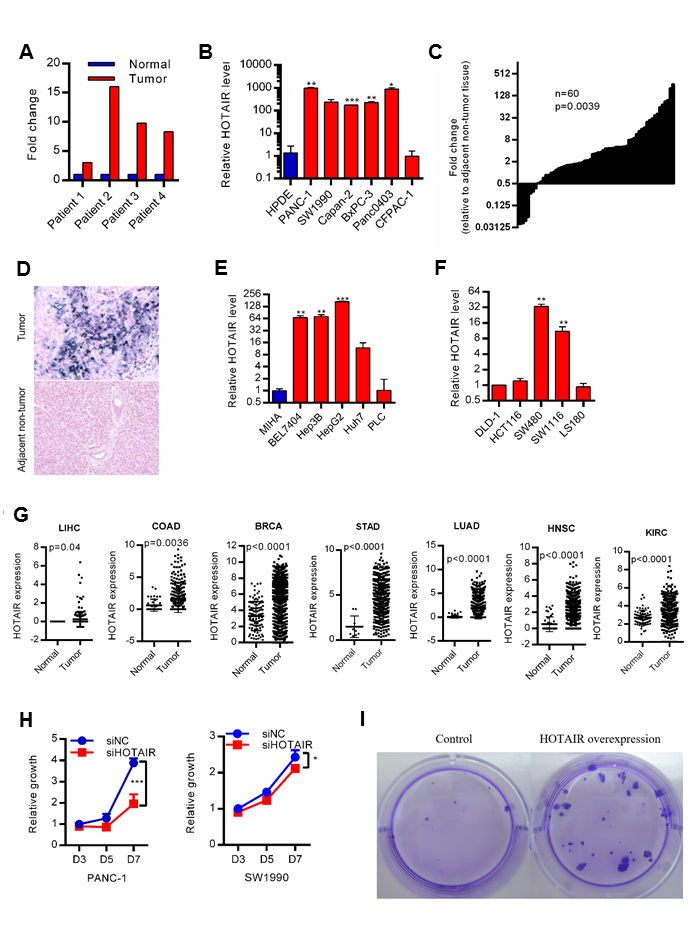

**Figure S1. HOTAIR is up-regulated in cancer and promotes PDAC carcinogenesis.** **(A)** Microarray analysis identified HOTAIR dysregulated in PDAC tumor, compared to adjacent non-tumor tissues. **(B-C)** qRT–PCR analysis of HOTAIR expression in pancreas ductal adenocarcinoma (PDAC) **(B)** cell lines and **(C)** tumors, *P* =0.0039, two-tailed paired Student’s t-test. Expression of HOTAIR in PDAC was compared to non-tumorigenic human pancreatic ductal epithelial (HPDE) cells or adjacent non-tumor tissues. **(D)** Representative In situ hybridization (ISH) images showing the expression of HOTAIR in PDAC tumor but not in adjacent non-tumor tissue. **(E-F)** HOTAIR expression in **(E)** hepatocellular carcinoma (HCC) and **(F)** colorectal carcinoma (CRC) cells. Expression of HOTAIR in HCC was compared to non-tumorigenic MIHA cells. **(G)** TCGA analysis of HOTAIR expression in LIHC, liver hpatocellular carcinoma; COAD, colorectal adenocarcinoma; BRCA, breast carcinoma; LUAD, lung adenocarcinoma; HNSC, head and neck squamous cell carcinoma. n = 2,816 samples. **(H)** cell proliferation assay after depletion of HOTAIR in PANC-1 and SW1990 cells. **(I)** Colony formation assay after overexpression of HOTAIR in HPDE cells. Data are from at least three independent experiments and plotted as means ± SD. * P < 0.05, * * P < 0.01,　* * * P < 0.001.

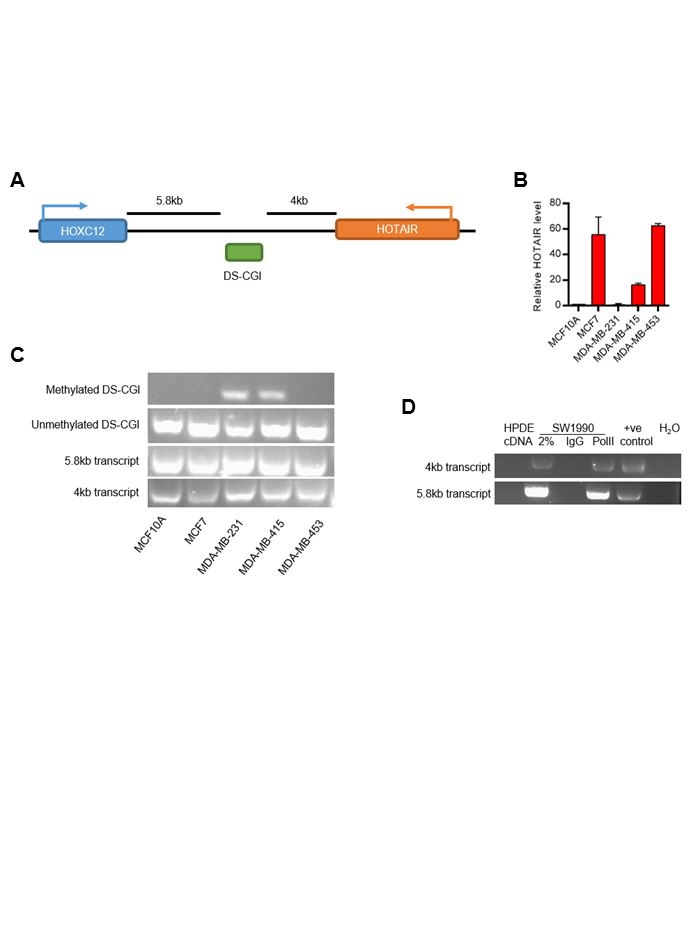

**Figure S2. Methylation of downstream CpG (DS-CGI) is not associated with HOTAIR expression. (A)** Location of DS-CGI island between HOTAIR and HOXC12. **(B)** qRT-PCR analysis of HOTAIR expression in breast cancer cells. Expression of HOTAIR in breast cancer was compared to non-tumorigenic human breast epithelial cells MCF10A. **(C)** Methylation status of DS-CGI and expression of both 5.8kb and 4kb transcripts in breast cancer cells and MCF10A cells. **(D)** Expression of both 5.8kb and 4kb transcripts in HPDE cells. ChIP analysis of RNA PolII occupancy on both 5.8kb and 4kb regions in SW1990 cells. Anti-H3 antibody was used as positive control. Data are from at least three independent experiments and plotted as means ± SD.

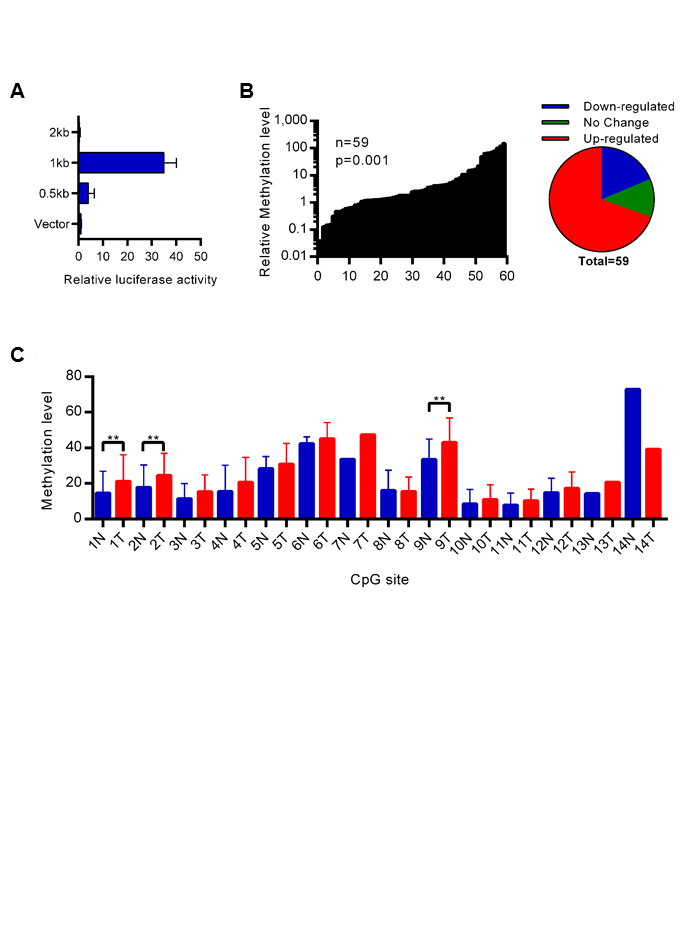

**Figure S3. Methylation of Ex-CGI is increased in PDAC tumors and is associated with HOTAIR expression. (A)** Luciferase assay to locate promoter of HOTAIR gene. **(B)** qMSP analysis of methylation level of Ex-CGI in PDAC tumors. **(C)** pyrosequencing analysis of methylation level of individual CpG site in Ex-CGI. Data are from at least three independent experiments and plotted as means ± SD.

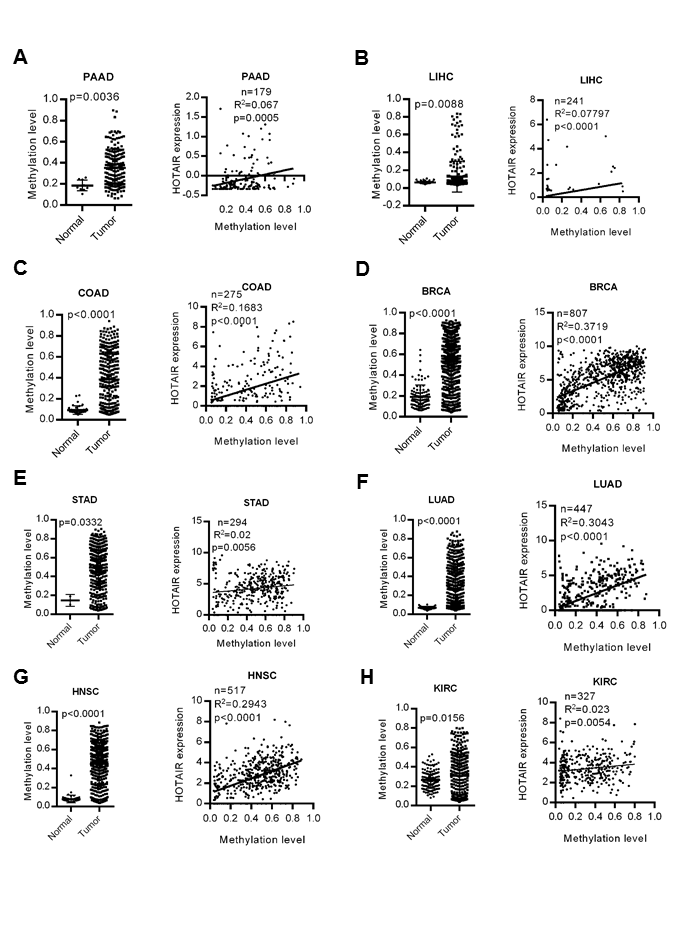

**Figure S4. Methylation of Ex-CGI is increased in cancers and is associated with HOTAIR expression.** TCGA analysis of Ex-CGI methylation level and its correlation with HOTAIR expression in **(A)** PAAD, pancreatic ductal adenocarcinoma; **(B)** LIHC, liver hpatocellular carcinoma; **(C)** COAD, colorectal adenocarcinoma; (**D**) BRCA, breast carcinoma; **(E)** LUAD, lung adenocarcinoma; and **(F)** HNSC, head and neck squamous cell carcinoma. n = 2,754 samples.

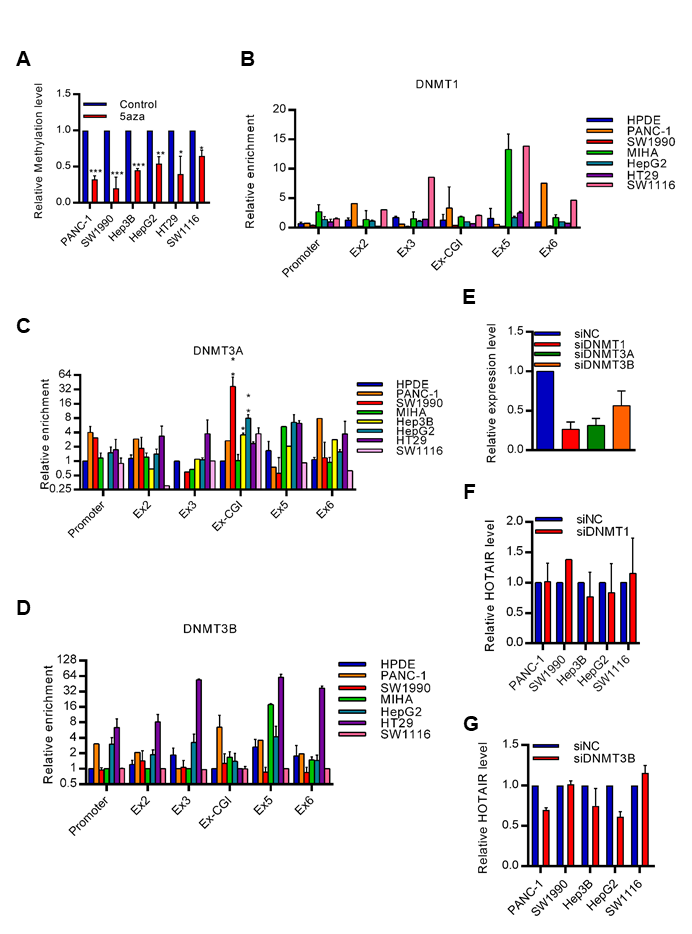

**Figure S5. Inhibition of Ex-CGI methylation inhibits HOTAIR expression.** **(A)** Ex-CGI methylation level after treatment with 5aza in PDAC, HCC and CRC cells. **(B-D)** ChIP analysis of **(B)** DNMT1, **(C)** DNMT3A and **(D)** DNMT3B on HOTAIR gene in PDAC, HCC and CRC cells. **(E)** Knock-down efficiency of siRNAs targeting DNMT1, DNMT3A and DNMT3B respectively. **(F-G)** HOTAIR expression after knock-down of **(F)** DNMT1 or **(G)** DNMT3B in PDAC, HCC and CRC cells. Data are from at least three independent experiments and plotted as means ± SD. * P < 0.05, * * P < 0.01,　* * * P < 0.001.

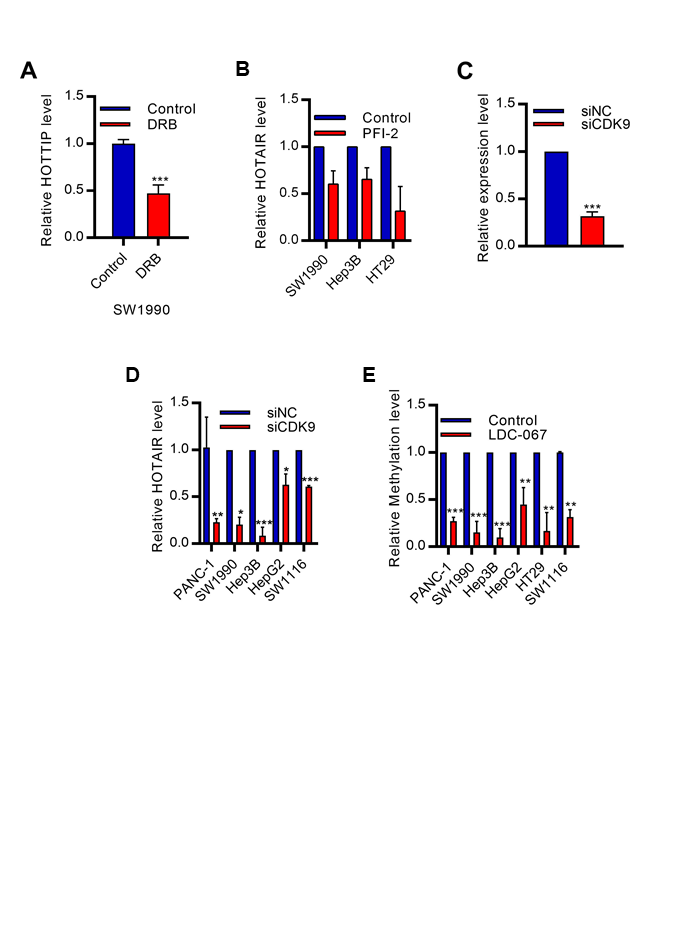

**Figure S6. Depletion of CDK7-CDK9 inhibits the expression of HOTAIR. (A-B)** HOTAIR expression level after inhibition of pTEFb by **(A)** DRB or **(B)** PFI-2 in PDAC, HCC and CRC cells. **c**, Knock-down efficiency of siRNAs targeting CDK9. **(D-E)** depletion of CDK9 by siRNA reduced **(D)** HOTAIR expression and **(E)** Ex-CGI methylation level in PDAC, HCC and CRC cells. Data are from at least three independent experiments and plotted as means ± SD. * P < 0.05, * * P < 0.01,　* * * P < 0.001

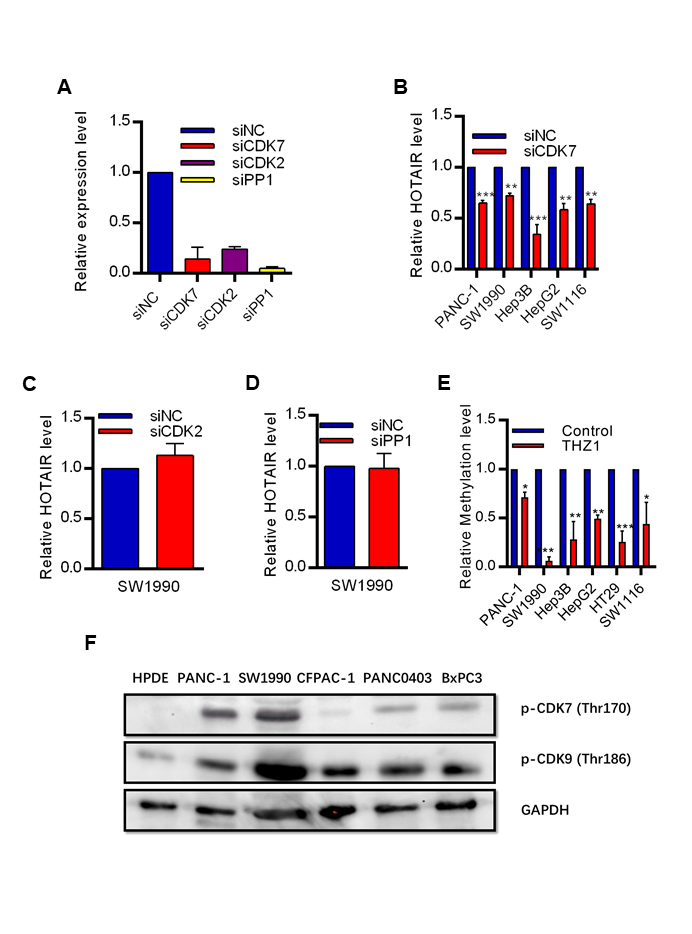

**Figure S7. CDK7 promotes transcription of HOTAIR in cancers.** **(A)** Knock-down efficiency of siRNA targeting CDK7, CDK2 and PP1 respectively. **(B-D)** HOTAIR expression after knock-down of **(B)** CDK7, **(C)** CDK2 and **(D)** PP1 by siRNAs. **(E)** depletion of CDK7 by siRNA reduced methylation level of Ex-CGI in PDAC, HCC and CRC cells. **(F)** Immunoblot analysis of CDK7 and CDK9 phosphorylation in PDAC cell lines. Data are from at least three independent experiments and plotted as means ± SD. * P < 0.05, * * P < 0.01,　* * * P < 0.001.

**Table S1. Primers used in this study**

| HOTAIR-qMSP-UF | TTAGGGATGAGTGATTGGATAGATTTG | qMSP |
| --- | --- | --- |
| HOTAIR-qMSP-UR | CCAACTAACTCAAACAACTCCCAC | qMSP |
| HOTAIR-qMSP-MF | GACGAGCGATTGGATAGATTCG | qMSP |
| HOTAIR-qMSP-MR | CGACTAACTCGAACAACTCCCG | qMSP |
| HOXC12-DS5.8K-qRTPCR-F | CACCCTCCCTGCTATATCCA | qRT-PCR |
| HOXC12-DS5.8K-qRTPCR-R | GACCAAAGGCTGAGCATAGG | qRT-PCR |
| HOXC12-DS4K-qRTPCR-F | AGGAGGTCTGTGCTTCAGGA | qRT-PCR |
| HOXC12-DS4K-qRTPCR-R | GTTCCAAGGGGTCTTTAGCC | qRT-PCR |
| HOTAIR qRTPCR F | ATCAGAAAGGTCCTGCTCC | qRT-PCR |
| HOTAIR qRTPCR R | GTCTGTAACTCTGGGCTCC | qRT-PCR |
| HOTAIR-qRTPCR-SF1 | GACAGGGTCTGGGACAGAAG | qRT-PCR |
| HOTAIR-qRTPCR-SR1 | GGAAATCAGGGCAGAATGTG | qRT-PCR |
| HOTAIR-qRTPCR-SF2 | GAGAGAGGGAGCCCAGAGTT | qRT-PCR |
| HOTAIR-qRTPCR-SR2 | TCAGACTCTTTGGGGCCTTA | qRT-PCR |
| HOTAIR-qRTPCR-SF3 | AATATCCCGGAGGTGCTCTC | qRT-PCR |
| HOTAIR-qRTPCR-SR3 | TCCCCTACTGCAGGCTTCTA | qRT-PCR |
| HOTAIR-qRTPCR-SF4 | GAGAGAGGGAGCCCAGAGTT | qRT-PCR |
| HOTAIR-qRTPCR-SR4 | TCAGACTCTTTGGGGCCTTA | qRT-PCR |
| HOTAIR-qRTPCR-SF5 | CTGGCAGAGAAAAGGCTGAA | qRT-PCR |
| HOTAIR-qRTPCR-SR5 | TACCAGGTCGGTACTGGCTTA | qRT-PCR |
| HOTAIR-ChIP-3'-P F | TCCCACCCCTCTCTTTTCTT | ChIP |
| HOTAIR-ChIP-3'-P R | TCGCGGCATTTTTATGAGAT | ChIP |
| HOTAIR-ChIP-5'-P F | CCATTTCCAGCCTGCTTTTA | ChIP |
| HOTAIR-ChIP-5'-P R | AGCAACCCGACCCTATTTCT | ChIP |
| HOTAIR-ChIP-3'-EX F | GGTCCTCCATTTCAGCCTTT | ChIP |
| HOTAIR-ChIP-3'-EX R | GAAGGAGGGGCGTCTTTATT | ChIP |
| HOTAIR-ChIP-5'-EX F | CGCCATGACAAAGTGAAGGT | ChIP |
| HOTAIR-ChIP-5'-EX R | AAGGCCCCAAAGAGTCTGAT | ChIP |
| HOTAIR-ChIP-Ex6-5'-F | TGGCCAAGCACCTCTATCTC | ChIP |
| HOTAIR-ChIP-Ex6-5'-R | GTGTAGACGCCGCCATATTT | ChIP |
| HOTAIR-ChIP-Ex2-F | AGAGAGCACCAGGCACTGAG | ChIP |
| HOTAIR-ChIP-Ex2-R | AGCACCTCCGGGATATTAGG | ChIP |
| HOTAIR-ChIP-Ex6-3'-F | CTAACTGGCAGCACAGAGCA | ChIP |
| HOTAIR-ChIP-Ex6-3'-R | GGGTCCCACTGCATAATCAC | ChIP |
| HOTAIR-ChIP-Ex3-F | CAGTGGAATGGAACGGATTT | ChIP |
| HOTAIR-ChIP-Ex3-R | CCCGCCTCTCTTTTTCTCTA | ChIP |
| HOTAIR-ChIP-Ex5-F | TTAATTCCCTCCTGCCCTTT | ChIP |
| HOTAIR-ChIP-Ex5-R | CCAGATAAGCCAGACCCAAT | ChIP |
| CDK7-F | TGAGGCGGGGATACTAAAGC | qRT-PCR |
| CDK7-R | TTCTCATAACGCTTTGCCCG | qRT-PCR |
| CDK9-F | GCATCATGGCAGAGATGTGG | qRT-PCR |
| CDK9-R | TTCAGCCTGTCCTTCACCTT | qRT-PCR |
| GAPDH-F | TGCCTCCTGCACCACCAACT | qRT-PCR |
| GAPDH-R | CCCGTTCAGCTCAGGGATGA | qRT-PCR |
